# The Inversion of Naïve-to-Memory T Cell Ratios With Age: Potential Links to Autoimmunity and the Role of Lifetime Infection History

**DOI:** 10.1101/2025.01.05.631399

**Authors:** Richard Murdoch Montgomery

## Abstract

As individuals grow older, their T cell repertoire progressively shifts from being dominated by naïve T lymphocytes to featuring a preponderance of memory T cells. This phenomenon—well-documented in immunological literature—is frequently attributed to repeated antigenic stimulation over the course of a lifetime, coupled with thymic involution and other age-associated changes. Although memory T cells are crucial for rapid responses to previously encountered pathogens, there is increasing interest in whether this dramatically shifted balance might predispose certain individuals to a higher risk of autoimmunity, especially when compounded by chronic stress and recurrent infections. In this article, we propose a conceptual model linking the age-related inversion of the naïve-to-memory T cell ratio to an elevated likelihood of immune dysregulation. We integrate mathematical equations to capture the dynamic interactions among key variables, present simulated data in four informative graphs, and discuss the implications and limitations of our approach. Our goal is to invite a deeper consideration of how infection history, stress-induced immune breakdowns, and the intrinsic remodeling of T cell populations with age could collectively contribute to autoimmunity.

## Section 1. Introduction

T cells are fundamental components of the adaptive immune system and are known for their extraordinary specificity in recognizing antigens. Their development occurs primarily in the thymus, where progenitor cells differentiate into naïve T cells that bear unique T cell receptors (TCRs) capable of binding specific peptides presented by major histocompatibility complex (MHC) molecules. In infants and young children, the thymus is relatively large and active. It produces a substantial output of newly formed naïve T cells, which remain largely unexperienced (i.e., they have not yet encountered their cognate antigen). *Studies consistently show that during early life, especially in infancy, more than 90%—some sources suggest up to 95%—of T lymphocytes in circulation can be classified as naïve [Britanova et al., 2016].* These cells feature a broad repertoire diversity, ready to mount primary immune responses against novel pathogens.

### Section 1. Aging

With aging, however, the composition of the T cell pool changes in a profound manner. Researchers have observed that memory T cells begin to dominate later in life, sometimes reaching proportions well above 80–90% in older adults [Nikolich-Žugich, 2018]. One of the major drivers of this phenomenon is the concept of thymic involution. Starting roughly in adolescence, the thymus undergoes a progressive reduction in size and functional capacity, leading to a markedly decreased output of recent thymic emigrants (RTEs). This reduction in naïve T cell production is further compounded by the repeated antigenic challenges individuals experience as they encounter various pathogens or pollution and chronic stress, both of which induce the clonal expansion of pathogen-specific T cells that, after clearing the infection, transition into memory cells. This shift in the naïve-to-memory T cell ratio is characteristic of the aging process and is often conceptualized under the broader umbrella of *“immunosenescence”* [Fulop et al., 2018]. Immunosenescence refers to the cumulative changes in the immune system that diminish its efficiency in mounting protective responses against new or evolving pathogens and, simultaneously, may predispose to a range of immunological dysregulations.

Aging-associated immunosenescence is not solely about the numeric decline of naïve T cells but also includes altered function of cells in both the innate and adaptive compartments*. Nonetheless, a central aspect of interest for clinicians and immunologists is understanding how a massively expanded population of memory T cells might affect immune homeostasis and, in certain contexts, facilitate the emergence of autoimmune pathologies.*Autoimmune diseases are complex disorders in which the immune system fails to maintain self-tolerance, resulting in a pathologic response directed against the body’s own cells and tissues. Examples range from systemic lupus erythematosus (SLE) to rheumatoid arthritis, type 1 diabetes, multiple sclerosis, fibromyalgia and others. Many of these conditions have multifactorial etiologies, encompassing genetic predisposition, hormonal influences, and a variety of environmental triggers [Abbas et al., 2018].

### Section 1.2 Possible Triggers

One longstanding hypothesis posits that infectious history may serve as a pivotal environmental determinant that influences the likelihood of autoimmunity. Certain infections can trigger immunological cross-reactivity, commonly referred to as “molecular mimicry,” where epitopes on a pathogen resemble self-antigens, thereby priming a T cell response that can inadvertently target host tissues [Amital et al., 2008]. Beyond specific molecular mimicry, the cumulative burden of infections can also remodel the T cell repertoire more globally. *Every infection recruits subsets of naïve T cells to expand and differentiate into effector cells, a fraction of which persist as memory T cells long after the pathogen is eliminated.*Over decades and through repeated infections, this memory subset inflates and may come to dominate the immune landscape. A consequence of this inflation is a reduced reservoir of truly naïve T cells capable of responding to new pathogens or playing a role in self-tolerance “education” [Zhang et al., 2019]. Additionally, older memory T cells may become functionally senescent, producing proinflammatory cytokines that can drive chronic low-grade inflammation. The term “inflammaging” has been introduced to describe the baseline inflammatory state often observed in older adults [Franceschi et al., 2018]. Chronic inflammation, in turn, can damage tissues, reveal cryptic antigens, and incite or exacerbate autoimmune processes.

It is also critical to acknowledge that not every individual with a high burden of infections develops an autoimmune disease, underscoring the complexity and multifactorial nature of immune tolerance. Genetic predisposition, sex hormones, and environmental factors such as the microbiome or exposure to toxins all modulate risk. Nonetheless, the correlation between extensive infection history and certain autoimmune conditions has been supported by epidemiological evidence. For example, some populations with high levels of infection have manifested unique patterns of immune reactivity, including expansions of particular T cell clones that may exhibit cross-reactivity to self-antigens [Dawes et al., 2019]. Stress, both chronic and acute, is another layer in this intricate puzzle. Psychological or physical stress has been shown to influence immune responses, altering the balance of T helper cell subsets, impacting regulatory T cell function, and potentially facilitating episodes of immune dysregulation through chronic elevated cortisol levels (Montgomery, 2024). Elevated cortisol levels can suppress certain arms of the immune response while promoting proinflammatory pathways in others, especially when stress is chronic rather than acute [Glaser and Kiecolt-Glaser, 2005]. This interplay among infection frequency, T cell repertoire shifts, and stress-induced vulnerabilities forms the crux of a “multiple hits” model, where repeated immune challenges and sporadic immunoregulatory breakdowns might tip the immune system toward pathogenic self-reactivity.

Investigations into the mechanisms behind T cell repertoire skewing and autoimmunity have yielded several points of interest. Some researchers emphasize the significance of T cell receptor (TCR) diversity. When naïve T cells are abundant, a wider range of TCR specificities is available, which can be beneficial not only for responding to novel pathogens but also for maintaining tolerance. Naïve T cells, having not yet encountered antigen, may be subject to ongoing peripheral tolerance checks, and some subset of them can be deleted if they exhibit dangerously high affinity for self-antigens. *With aging and with repeated expansions of memory T cells, the overall repertoire can become more oligoclonal or less diverse, thus reducing the ability of the immune system to appropriately respond to new antigens or to keep in check any reactivity that spontaneously arises against self.* Another avenue of investigation focuses on regulatory T cells (Tregs), which act as “peacekeepers” that quell excessive immune responses, including those that are self-reactive. Tregs also exhibit functional changes with aging; some evidence suggests that older individuals show a reduction in Treg efficacy or number, weakening the final line of defense against autoimmune reactions [Abbas et al., 2018]. On the other hand, certain data indicate that Tregs can sometimes increase with age but may lose functional capacity. Clearly, the story is far from simple and is influenced by multiple mediators.

### Section 1.3. The Role of Stress

Given these biological underpinnings, one can see how stress events, which can modulate Treg function or induce systemic proinflammatory states, might act as catalysts for autoimmune flares. Stress is not only psychological but may also arise from physical or metabolic sources, including surgery, trauma, malnutrition, or concurrent chronic diseases*. All of these insults can transiently disable or distract regulatory components, creating windows during which self-reactive T cells, if present in the memory pool, could expand.*The synergy between stressors and repeated infections, therefore, becomes particularly important. In societies or environments where infectious diseases are frequent, individuals may accumulate large expansions of memory T cells with various specificities. When stress or immunoregulatory breakdown occurs, some of these memory clones may become inadvertently activated, unleashing pathological responses. The question remains whether this repeated cycle of infection-driven memory T cell expansion and stress-induced regulatory lapses can be mathematically captured or predicted in a manner that can help guide clinical or public health interventions.

### Section 1.4 Mathematical Modeling

In immunology, mathematical modeling has a long and rich tradition. Classic models of infection dynamics, such as those pioneered by Anderson and May [Anderson and May, 1991], have expanded into more complex frameworks that can incorporate the nuances of adaptive immunity. Models aiming to capture T cell dynamics frequently employ systems of differential equations, tracking how different T cell subsets (e.g., naïve, effector, memory, regulatory) evolve over time. *These models can be extended to include the effects of stress, infection rates, and even the stochastics of antigenic exposure.*The conceptual approach is typically to specify rates of production, proliferation, and death for each subset, along with transition terms that reflect how naïve T cells become effector or memory T cells upon stimulation. By adjusting parameters to represent either high or low infection burdens, variable stress episodes, or different baseline rates of thymic involution, one can simulate distinct immunological trajectories and compare them with empirical data.

Empirical data for validating such models often come from longitudinal cohort studies that measure T cell subsets in participants over several years. These can involve enumerating naïve and memory T cells via flow cytometry, sequencing TCR repertoires to assess diversity, and evaluating clinical outcomes in terms of infection frequency and autoimmune markers. Although no single dataset is likely to provide a definitive test of all aspects of a model, integrated analyses that combine observational data from epidemiological studies, mechanistic insights from smaller immunological experiments, and well-designed computational models can together yield a coherent picture of how repeated immunological assaults shape T cell populations (a study like this is very likely to be tailored by advanced AI, not so far from the present).

One of the remarkable aspects of the human immune system is its capacity for memory, which is evolutionarily advantageous in safeguarding the host against recurrent pathogens. Yet, the very feature that grants this protective capacity might, under certain conditions, predispose the system to autoreactivity and chronic inflammation and frailty (Montgomery, 2024a). The question of how often and to what extent the normal, beneficial memory response tips over into a pathological autoimmune response remains an area of active debate. There is, however, an increasing consensus that factors like advanced age, high cumulative infection load, and recurrent stress episodes likely increase the probability of crossing this threshold. *Vaccination rates, in this context, play an important role of protecting against recurrent infections*.

This article aims to synthesize these ideas into a cohesive narrative: the inversion of naïve-to-memory T cells over time, exacerbated by multiple infections and stress events, may have a tangible association with the rising prevalence of autoimmune disorders in older populations or in individuals with particularly burdened immunological histories. We propose a simple set of ordinary differential equations (ODEs) that encapsulate the essential components of naïve T cell decline, memory T cell accumulation, and superimposed episodes of increased antigenic stimulation and stress-induced dysregulation. We also present a series of simulations that illustrate how different parameter settings— such as higher infection frequency or additional stress events—can accelerate the shift toward memory T cell dominance and potentially push an individual’s immune system toward an *“autoimmune threshold.”*

By articulating these theoretical relationships in a quantitative framework, researchers and clinicians can begin to ask more precise questions about interventions. For example, would reducing the burden of certain persistent infections (like cytomegalovirus, which is associated with “memory inflation” [Karrer et al., 2003]) preserve a healthier naïve T cell pool and thereby mitigate autoimmune risk? Could interventions aimed at stress reduction—through psychological counseling, lifestyle modifications, or targeted medical therapies— help bolster regulatory T cell function or otherwise buffer the immune system from tipping into pathological autoreactivity? Furthermore, could early vaccination strategies that prime broad protective immunity reduce the cumulative immune burden that arises from repeated, uncontrolled infections, thereby preserving a more balanced T cell repertoire?

Indeed, each of these questions forms part of a broader research agenda that seeks to unravel the relation between infection, stress, aging, and autoimmunity. The notion that older adults frequently exhibit compromised responses to novel infections (as seen with influenza or emerging pathogens) while showing increased prevalence of autoimmune phenomena highlights the dual nature of an immune system dominated by memory T cells. On one hand, these memory T cells deliver rapid recall responses to previously encountered pathogens. On the other hand, they may hamper the immune system’s flexibility and tolerance mechanisms. Since many autoimmune conditions are polygenic and exhibit significant variability in onset and severity, population-level analyses will inevitably yield heterogeneous outcomes*. Nevertheless, the robust correlation between advanced age, decreased naïve T cell numbers, heightened memryoT cell dominance, and autoimmune conditions across a range of demographics supports the line of reasoning presented here [Carr et al., 2022]*.

To conclude, the introduction sets the stage for our central premise: that the age-related shift from a naïve T cell-dominated repertoire to a memory T cell-dominated repertoire can be accelerated by repeated infections and episodes of stress, and that this acceleration correlates with an elevated risk of autoimmunity. The article proceeds by outlining a mathematical model that captures these dynamics in an accessible way. We then present simulation results—displayed in four graphs—that highlight how different parameters can reshape T cell populations over time. Finally, we grub around into a discussion that interprets these findings in the broader context of immunological theory and clinical evidence, addressing the strengths, limitations, and future prospects of this approach.

## Section 2. Methodology

### Section 2.1 Model Construction

To study how repeated infection episodes and stress events might drive the inversion of naïve-to memory T cell ratios and subsequently influence autoimmunity risk, we constructed a system of ordinary differential equations (ODEs). We define:

- *N*(*t*) : The population of naïve T cells at time *t*.
- *M*(*t*) : The population of memory T cells at time *t*.
- *a* : The baseline thymic output of naïve T cells.
- *β* : A constant describing the rate of decline in thymic output due to involution.
- *γ* : The rate constant for the conversion of naïve *T* cells to memory *T* cells upon antigenic stimulation.
- *δ_N_* : The turnover (death) rate of naïve T cells.
- *δ_M_* : The turnover (death) rate of memory T cells.
- *f*(*t*) : A function describing the burden of infection over time (i.e., how often and how intensely new antigenic exposures occur).

We assume the total T cell population is governed by the following set of ODEs:

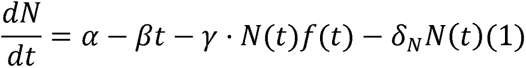

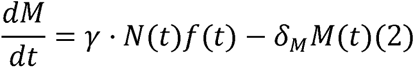

The term *α*-*βt* models a linearly decreasing thymic output as a proxy for thymic involution over time. The product γ. *N*(*t*)*f*(*t*) represents the stimulation-driven conversion of naïve *T* cells into memory T cells, which we assume depends on both the magnitude of the naïve T cell pool and the infection function *f*(*t*). Turnover (death or functional dropout) of naïve and memory cells is represented by *δ_N_N*(*t*) and *δ_M_M*(*t*), respectively.

### Section 2.2. Modeling Infection Burden and Stress

In order to capture episodic infections, we let *f*(*t*) be a sum of Gaussian functions. Each Gaussian represents a distinct infection event with a specified amplitude, timing, and duration:

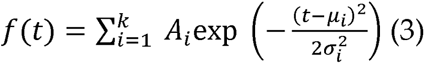

where k is the total number of significant infection episodes over the lifespan, *A_i_* is the peak magnitude of infection *i*, *µ_i_* is the time at which infection *i*, peaks, and *σ_i_* determines the width or duration of that infection peak.

Stress events are modeled by allowing transient increases in the conversion rate γ. Specifically, we define a small "stress factor" *η*(*t*) that becomes positive during stress episodes, which effectively modifies γ into γ. (1 + *η*(t)). This approach ensures that during stressful periods, the naïve T cells exposed to antigens have a higher likelihood of converting into memory T cells.

## Section 4. Results

We implemented the equations described above in Python (see Attachment). The simulation proceeds via a simple Euler method, and we track the populations and over the course of an 80-year lifespan. Four representative graphs illustrate different scenarios:

### 1. Figure 1-Baseline Naïve and Memory Dynamics

Shows how cells gradually decrease while memory cells steadily increase under moderate infection rates and periodic stress events.

**Figure 1.**
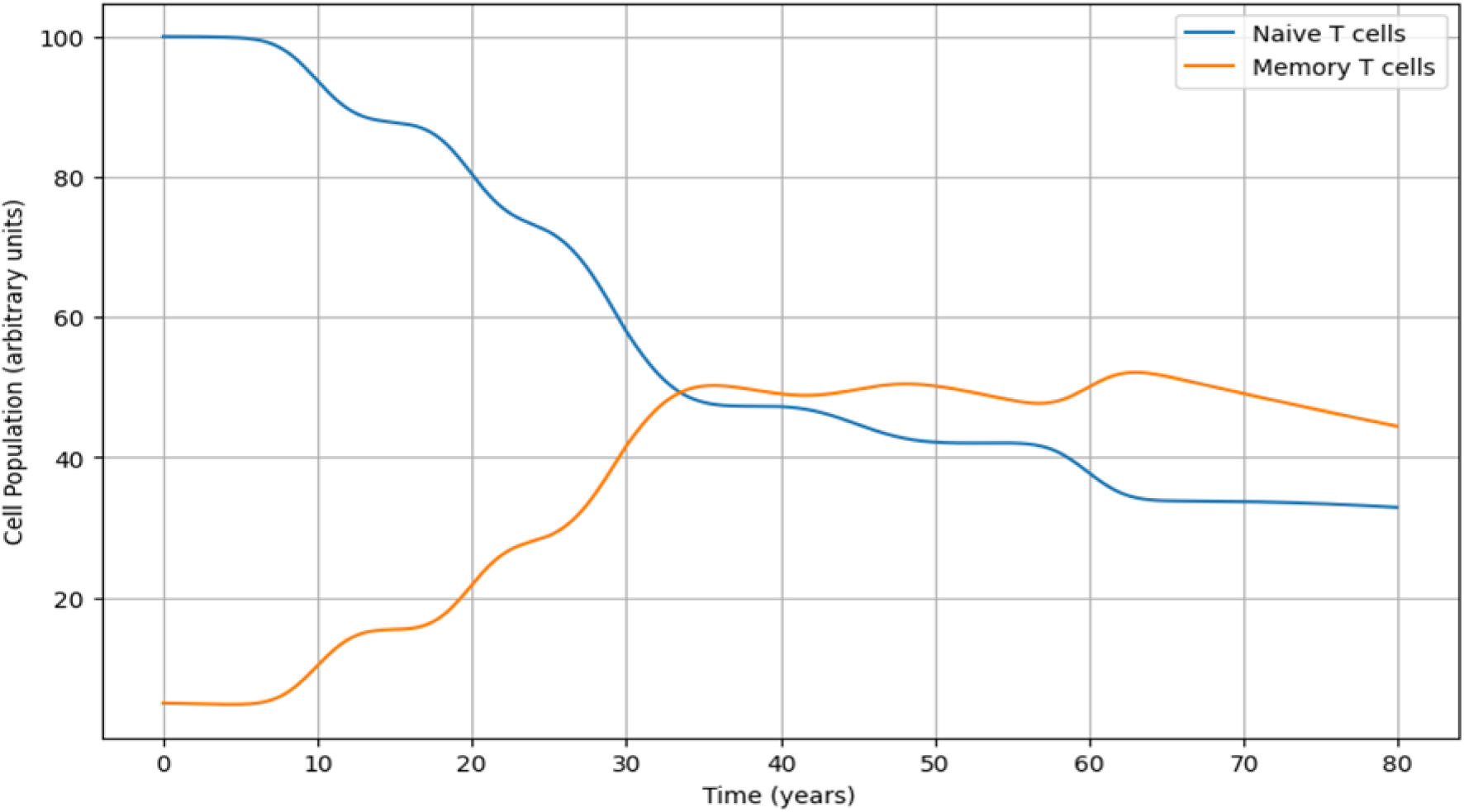
Baseline Naïve and Memory Dynamics.

### 2. Figure - High Infection Burden

Demonstrates how frequent and intense infection events accelerate the decline of cells and promote an earlier and more pronounced memory-dominated landscape.

**Figure.**
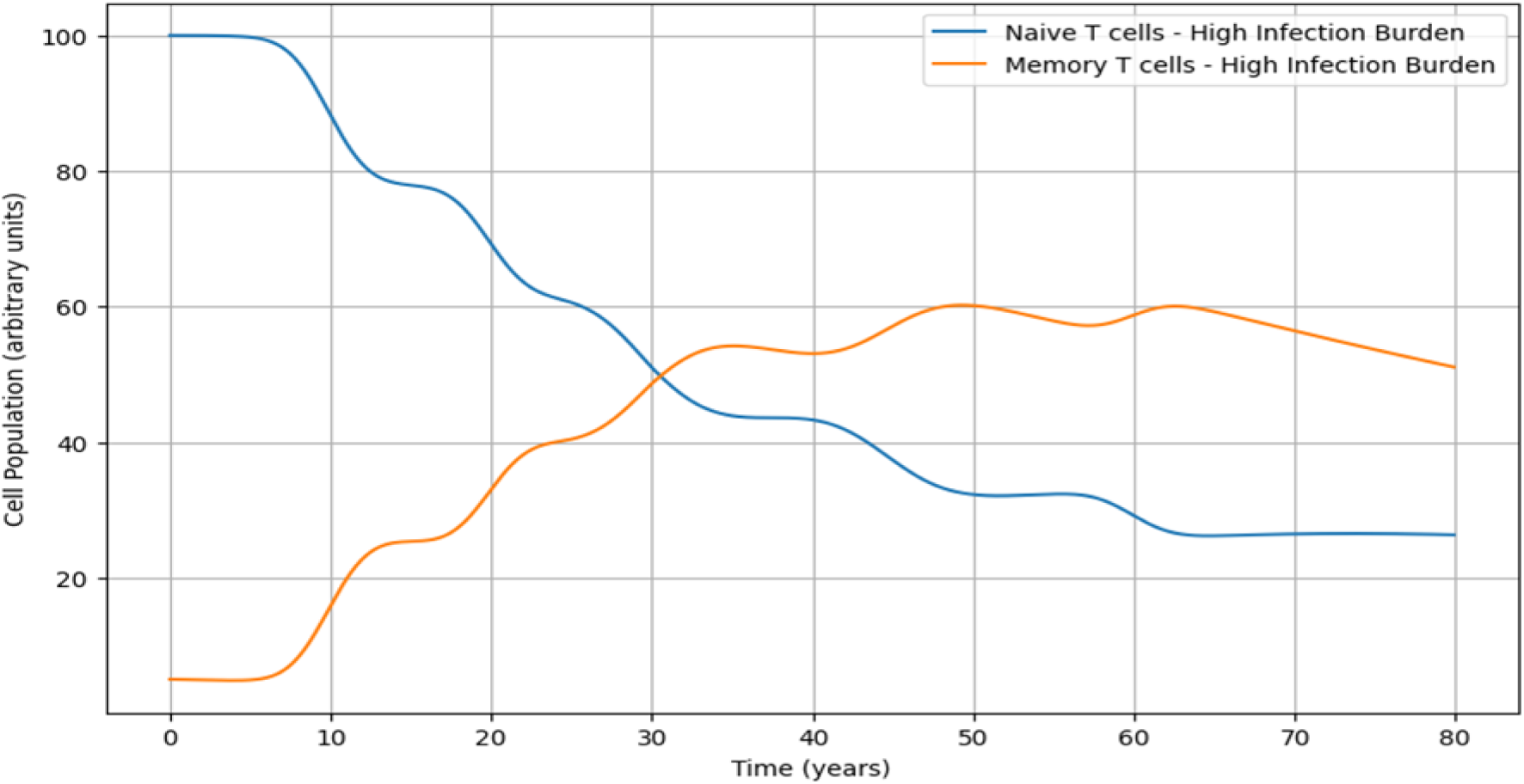
High Infection Burden.

### 3. Figure 3. - Impact of Stress Events

Explores how superimposing stress episodes amplifies the rate of *Figure 4 - Potential Autoimmune Threshold*.

**Figure 3.**
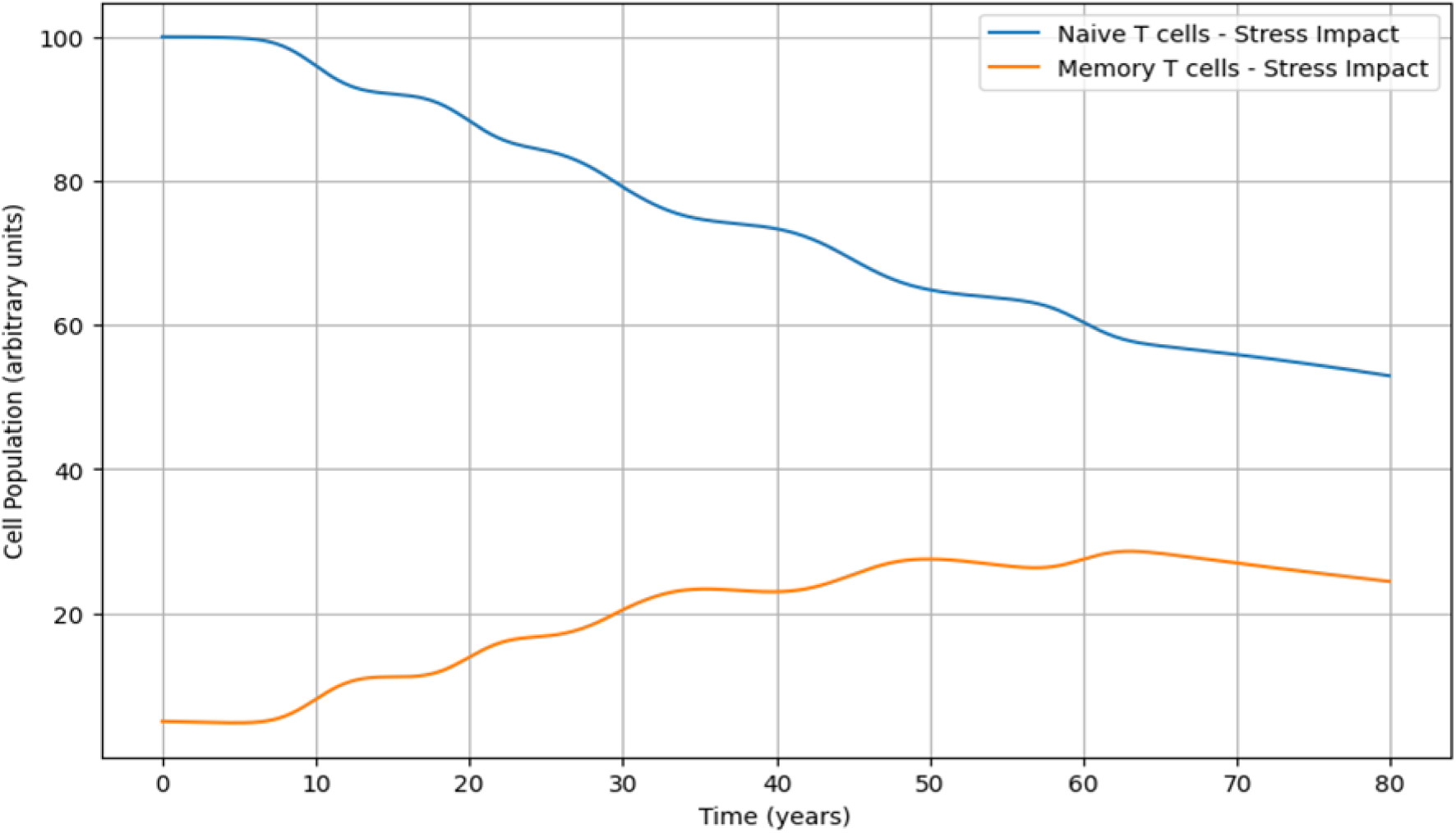
Impact of Stress Events.

**Figure 4.**
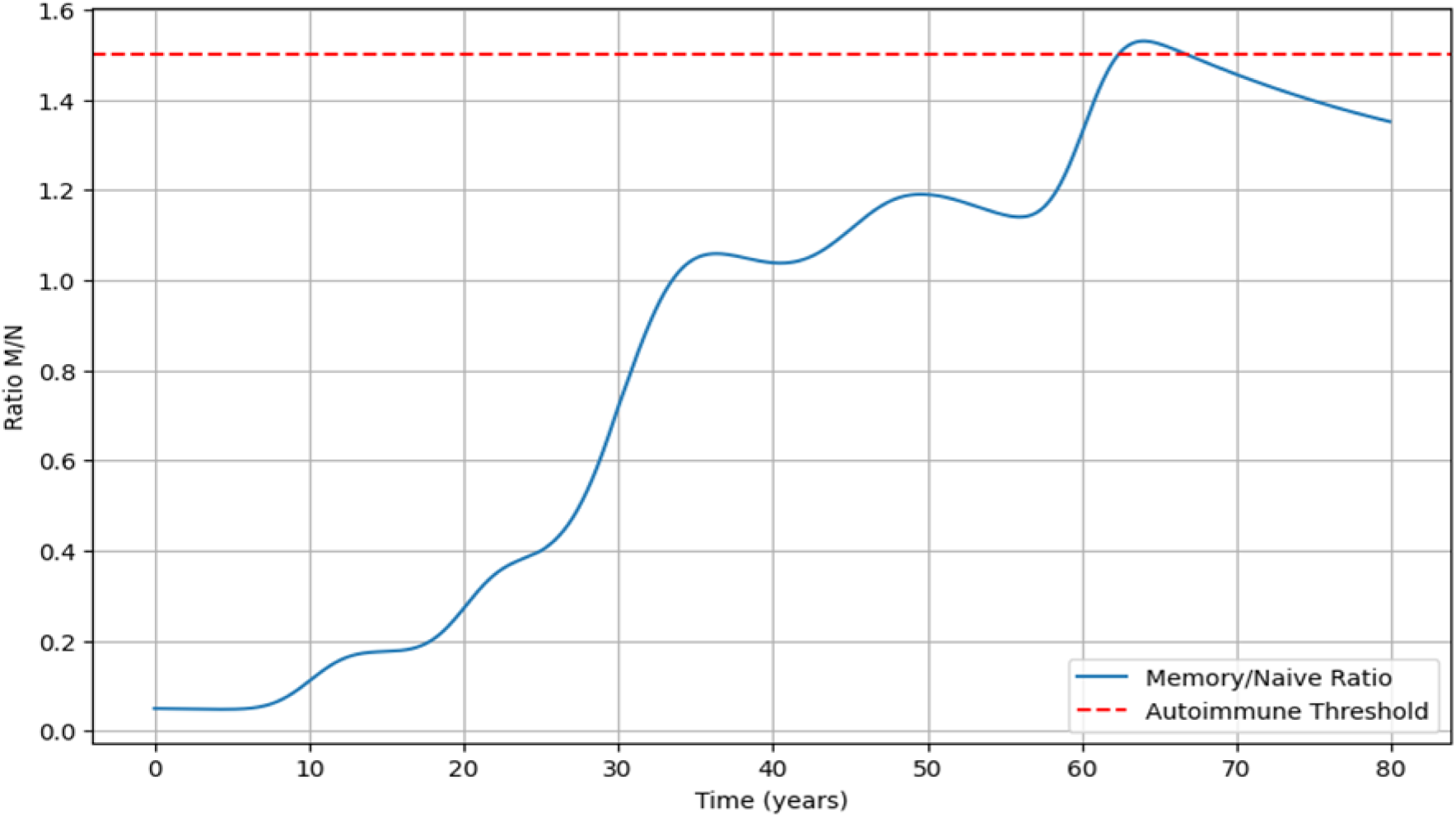
Potential Autoimmune Threshold.

-to-memory conversion, leading to a steeper reduction in naïve *T* cells and a faster increase in memory *T* cells.

### 4. Figure 4 - Potential Autoimmune Threshold

Depicts a conceptual threshold based on the ratio of memory to *naïve* T cells, suggesting that once this ratio passes a certain point, the probability of autoreactivity may rise dramatically.

These figures collectively reinforce the idea that both infection frequency and stress can shape the age-related transition from a *naïve* -rich T cell pool to a memory-dominated one. In the next section, we examine these findings in the context of existing immunological knowledge and address the strengths, weaknesses, and broader implications of our modeling approach.

## Section 4. Discussion

The simulation results presented herein prompt deeper reflection on how repeated infections, stress episodes, and a baseline age-driven decline in naïve T cell output might collectively promote or accelerate the skewing of the T cell repertoire toward memory dominance. This skewing carries significant implications for autoimmunity, as expanded memory populations can potentially harbor clones primed for self-reactivity. A crucial point to consider is the interplay of biological processes that gradually reduce thymic function over time while simultaneously favoring the survival and persistence of memory T cells, especially among those T cells that have encountered common pathogens. It is well established in immunology that thymic involution begins in adolescence [Nikolich-Žugich, 2018], although the precise rate and extent of this decline varies considerably among individuals, partly due to genetic and environmental influences [Carr et al., 2022]. Our model, which uses a simple linear term (*α* - *βt*) to approximate thymic output, is intentionally simplified; in reality, thymic involution can follow more intricate trajectories, sometimes with periods of relative stability interspersed with sharper declines, potentially influenced by hormonal changes or acute health events [Fulop et al., 2018]. Nevertheless, even this simplified representation captures the well documented phenomenon that the capacity to generate new naïve T cells diminishes with advancing age [Abbas et al., 2018].

Over the last few decades, investigators have shown that elderly individuals not only have fewer naïve *T* cells but also display expansions of memory *T* cell clones specific to pathogens such as cytomegalovirus (CMV). This process, frequently referred to as "memory inflation", becomes increasingly evident as repeated infections stimulate the continual recruitment of naïve T cells and their subsequent differentiation into long-lived memory populations [Karrer et al., 2003]. Once formed, many of these memory *T* cells can persist for extended durations- some subsets for years or even decades-particularly if low-level antigenic stimulation drives their maintenance and slow proliferation [Zhang et al., 2019]. This adaptive trait ensures robust immunological memory but also carries a cost: fewer resources (e.g., cytokines, growth factors, and suitable niches) remain available for naïve T cells, and the generation of new clones essential for responding to novel pathogens or for maintaining an extensive, self-tolerant repertoire becomes more constrained [Britanova et al., 2016]. In our model, the infection function *f*(*t*), composed of Gaussian pulses, illustrates how discrete infection events incrementally tilt the immune system toward memory dominance (for example, in big cities with a high pollution rate). While real-world infections may rarely follow neat Gaussian distributions - coinfections, subclinical infections, and vaccinations frequently overlap in complex ways - the fundamental principle persists that each immunological challenge can leave behind an expanded contingent of memory cells, especially if the infection triggers large-scale T cell clonal expansion.

### Section 4.1 The Stress Factor

Superimposing stress episodes upon this phenomenon, as our model demonstrates, can further accentuate the shift. *Stress is well known for reshaping the immune landscape via alterations in cortisol levels, shifts in cytokine profiles, changes in metabolism, and fluctuations in sympathetic nervous system activity [Glaser and Kiecolt-Glaser, 2005].* Depending on context, stress may induce a state of generalized immune suppression-impairing certain T cell subsets or their functional capacities—or foster a proinflammatory milieu that favors memory T cell activation at the expense of effective regulatory cell function [Amital et al., 2008]. Our simulated "stress factor" *η*(t), which transiently boosts the conversion rate r, is an abstraction of these complex neuroendocrine-immune interactions. Even modest, short-lived increases in the naïve-to-memory transition rate, *if they occur recurrently*, accumulate over time, compounding naïve T cell depletion and driving the proliferation of memory clones.

### Section 4.2. Broader Implications

The broader implications of these observations are significant. Individuals who experience multiple infections over their lifetime or who endure sustained or repeated stress exposures might show a particularly swift or dramatic inversion in their naïve-to-memory T cell ratio [Zhang et al., 2019]. *An elevated memory T cell compartment, especially one enriched in senescent or "exhausted"T cells, is more likely to generate high levels of proinflammatory mediators such as interleukin-6 (IL-6), tumor necrosis factor-alpha (TNF-* a *), and interferon-gamma (IFN-* r *) [Franceschi et al., 2018].* This chronic, low-grade inflammatory background can damage tissues, exposing cryptic or sequestered antigens. Once revealed, such antigens might be erroneously recognized by autoreactive T cell clones persisting in the memory pool due to incomplete central (thymic) or peripheral tolerance. Chronic inflammation itself can undermine the normal function of regulatory T cells (Tregs), thereby diminishing their capacity to restrain self-reactive clones [Abbas et al., 2018]. In turn, this sets in motion a vicious cycle: frequent infections expand memory *T* cells, some of which may harbor autoreactive potential; these memory T cells drive inflammation and tissue damage; new antigens become accessible; and compromised Treg function fails to contain the subsequent expansion of self-targeting clones [Fulop et al., 2018].

### Section 4.3. Autoimmune Diseases

From a clinical vantage, numerous studies underscore connections between chronic infections and certain autoimmune diseases. For instance, Epstein-Barr virus (EBV) infection has been linked epidemiologically to an increased risk of developing multiple sclerosis, even though the precise mechanism (possibly involving molecular mimicry or bystander activation) remains under active investigation [Zhang et al., 2019]. Hepatitis C virus infection has been implicated in mixed cryoglobulinemia and autoimmune-like rheumatologic conditions, while Helicobacter pylori infection has shown variable associations with immune-mediated disorders such as idiopathic thrombocytopenic purpura (ITP). Although some of these examples might involve direct molecular mimicry-where pathogen-derived epitopes resemble self-antigens-many also broadly illustrate how repeated or persistent antigenic stimulation can derail immune tolerance [Abbas et al., 2018]. Our model does not delve into the molecular details of antigen mimicry or bystander activation, but it supplies a theoretical framework for understanding how frequent immunological assaults can incrementally shift equilibrium away from a naïve-dominant state.

Quantifying "autoimmune risk" in a numerical simulation remains a formidable challenge. Our use of a conceptual "autoimmune threshold" superimposed on the memory-to-naïve *T* cell ratio is undeniably simplified, but this idea resonates with existing immunological discussions about tipping points in immune homeostasis [Dawes et al., 2019]. Past a certain ratio-or once a certain threshold of chronic inflammation and Treg impairment is reached - clinical autoimmunity might manifest. Observationally, many autoimmune diseases do increase in incidence with age, though the onset can differ considerably by condition; certain diseases, such as type 1 diabetes, can occur earlier in life, indicating that genetic predispositions, environmental triggers, and immunological complexity must be carefully dissected [Abbas et al., 2018]. Still, the broader trend of older populations showing greater prevalence of autoimmune disorders is consistent with an immune system that becomes progressively less tolerant and more prone to erroneous memory expansions.

### Section 4.4. Limitations

From the standpoint of our modeling framework, it is crucial to underscore the simplifications involved. Our equations focus principally on *CD4* ^+^*T* cell subsets (naïve vs. memory) and omit *CD8* ^+^ T cells, B cells, innate immune cells, or specialized subpopulations like Tregs, T follicular helper cells (Tfh), and Th17 cells. Each subset possesses distinct homeostatic dynamics, cytokine signatures, and roles in immune tolerance or pathology, making them essential for a nuanced comprehension of autoimmunity [Murphy and Weaver, 2017]. Moreover, our constant death rates a_N_ and a_M_ and a fixed conversion rate r do not wholly capture the complexities of aging biology, since turnover and responsiveness can alter dramatically with age or disease states [Zhang et al., 2019]. Our stress function *η*(t) is likewise a coarse stand-in for a kaleidoscope of neuroendocrine-immune interactions that vary among individuals. Stress is rarely uniform in duration or intensity, and psychological, physical, or metabolic stressors can impart distinct immunological signatures [Glaser and Kiecolt-Glaser, 2005].

Despite these recognized limitations, an ODE-based model offers a powerful lens for grasping fundamental population-level principles and systematically probing how different parameters shape outcomes. By generating scenarios in which infection is frequent, and stress levels are high, we can watch the immune system pivot more quickly toward a memory-dominated state, a phenomenon reflected in certain high-burden populations. Conversely, less frequent infection or effectively managed stress theoretically preserves a greater fraction of naïve *T* cells well into adulthood, at least within our simulations. The resonance of these patterns with real-world observations among older adults burdened by recurrent infections or chronic stress lends credibility to the conceptual underpinnings of the model, even though the latter cannot replicate all the fine-grained realities of human immunity [Franceschi et al., 2018].

Expanding our scope, it is important to remember that T cells rarely operate in isolation. Autoimmune diseases typically involve intricate crosstalk among T cells, autoreactive B cells, local tissues-resident immune cells, inflammatory mediators, and various tissue-specific factors [Abbas et al., 2018]. For example, T cells infiltrating the joints in rheumatoid arthritis, the central nervous system in multiple sclerosis, or the pancreatic islets in type 1 diabetes encounter local microenvironments shaped by chemokine gradients, endothelial adhesion molecules, and tissue resident antigen-presenting cells. Incorporating these local contexts into a broader model remains a substantial undertaking, yet the essential query of how an overabundance of memory T cells - many potentially senescent or dysfunctional - drives a proinflammatory milieu is pivotal to disentangling the roots of autoimmunity [Fulop et al., 2018].

Further complicating matters is the reliability-but also potential hazard-of memory T cells as the body’s immunological archives. Memory T cells not only rise in number with repeated infections but sometimes undergo phenotypic shifts, such as the appearance of the TEMRA subset (CD45RA reexpressing effector memory T cells), which can accumulate with age [Zhang et al., 2019]. These cells often produce copious proinflammatory cytokines and exhibit potent cytotoxic capabilities, potentially amplifying tissue damage if they become activated against self-antigens. *While memory T cells are crucial for rapid protection against familiar pathogens, they can become detrimental under conditions of chronic inflammation or repeated stress.*Thus, a memory-dominant immune system, though advantageous for recall immunity, can become maladaptive if regulation falters or if senescent clones proliferate unchecked [Nikolich-Žugich, 2018].

*Socioeconomic and epidemiological dimensions must also be considered. In communities or nations with limited healthcare resources, higher burdens of chronic infectious disease, and prevalent socioeconomic stressors, the shift toward a memory-dominant T cell profile may occur earlier or more intensively than in populations enjoying robust public health measures and lower infection rates [Carr et al., 2022].* This differential immunological aging could, in principle, affect the incidence and distribution of autoimmune diseases worldwide. Still, paradoxically, autoimmune disorders can also be more frequently diagnosed in resource-rich settings where advanced medical diagnostics are available. Hence, thorough epidemiological studies must account for not only infection quantity but infection quality (i.e., the nature of pathogens), nutritional status, psychosocial stressors, and genetic susceptibility. The interplay of these factors is extraordinarily intricate, but the integrative framework proposed in our model can help conceptualize how repeated immune challenges influence lifelong immune architecture and shift the balance toward pathological self-reactivity.

Ultimately, the thrust of our findings is that frequent or intense infection events, paired with stress induced increases in naïve-to-memory T cell conversion, may expedite the well-known age-related inversion of T cell subsets [Zhang et al., 2019]. Our model offers an initial scaffold for immunologists and clinicians to explore mechanistic links between memory T cell inflation, chronic inflammation, and heightened autoimmune risk. While stress effects are fluid and autoimmunity is multifactorial, the concurrence of immunosenescence, recurrent infection, and sporadic immune dysregulation stands as a plausible explanatory thread weaving through epidemiological data and clinical observations [Glaser and Kiecolt-Glaser, 2005; Britanova et al., 2016]. This integrated perspective has relevance for developing diagnostic tools to monitor vulnerable populations, preventive strategies to mitigate harmful recurrent infections or excessive stress, and therapeutic avenues that selectively target memory T cells bearing pathological potential. Although no model can incorporate all biological layers, the alignment of our results with established knowledge about T cell aging and infection-driven immunomodulation affirms that these processes are interconnected.

### Section 4.5 Perspectives

Going forward, refining this model to incorporate additional immune subsets (CD8 ^+^*T* cells, Tregs, innate lymphoid cells), feedback loops involving cytokines, and more nuanced representations of chronic infection could yield deeper insights. Furthermore, future work could integrate real-world longitudinal data on infection histories, stress biomarkers, and T cell subset profiling to calibrate our parameters more accurately and identify which individuals are most at risk for autoimmunity. Such endeavors may pave the way for personalized interventions that account for both genetic and environmental factors. In doing so, we move closer to an era of precision immunology, where preventing or delaying autoimmune conditions becomes an achievable aim rather than a distant aspiration.

In sum, this discussion underscores a central message: repeated immunological assaults-be they infections or stress-related-can accelerate the shift from a naïve-rich to a memory-rich *T* cell landscape, and such shifts correlate with increased autoimmune vulnerability in genetically or environmentally predisposed individuals. The synergy between infection burden, stress, and aging complicates the immunological balance but also presents opportunities for clinical intervention. By continuing to refine computational models and integrate them with epidemiological and integrate them with epidemiological and experimental data, researchers can better predict when and how the immune system might falter, ultimately guiding strategies that preserve immune tolerance and foster healthy aging.

## Section 5. Conclusion

The central premise of this work has been to illustrate how the human immune system, over a lifetime, transitions from a predominantly naïve T cell landscape to one replete with memory T cells, and how repeated infection episodes and periods of stress can hasten this transformation. In early life, the thymus is a robust organ that continually supplies fresh naïve T cells, each with a unique receptor repertoire ready to respond to novel antigens. However, thymic involution—commencing from late adolescence onward—results in a steady decline in the output of new T cells. When superimposed with recurrent microbial infections, immunological challenges, and stress-induced dysregulation, the system may reach a tipping point more quickly, shifting into an immunological state characterized by a large proportion of memory T cells and a diminished pool of naïve T cells.

*One of the salient considerations raised is the potential association between this memory-dominated T cell profile and an increased susceptibility to autoimmune disorders*.

Although memory T cells provide critical “immunological recall” and rapid pathogen clearance, there is growing evidence that a heavily skewed memory repertoire may also invite immune dysregulation, particularly in genetically susceptible individuals or in those experiencing chronic or repeated bouts of stress. The expanded memory compartment, in the context of reduced thymic output, may limit the capacity of the immune system to replenish the naïve T cell pool. This reduced naïve population is less adaptable to newly emerging pathogens and less capable of maintaining a broad, flexible repertoire that can “police” autoreactive clones. *Furthermore, senescent or functionally “exhausted” memory T cells can contribute to a background of chronic low-grade inflammation— often termed “inflammaging”—which may damage tissues and expose cryptic self-antigens, feeding into a cycle that fosters or exacerbates autoimmune phenomena*.

By applying mathematical models, we gain a clearer picture of how rates of infection, stress-induced immune modifications, and thymic involution come together to shape T cell dynamics. Although our model is simplified, it points to key parameters—such as infection frequency, stress intensities, and thymic output rates—that could be experimentally monitored and potentially targeted in interventional strategies. There is considerable scope to refine the equations used here by incorporating additional immune subsets and biological processes, most notably regulatory T cells (Tregs), which are crucial for maintaining self-tolerance. Tregs could provide a moderating influence in the model, offsetting the proinflammatory and potentially autoreactive tendencies of a memory-heavy repertoire. Chronic infections, too, might be modeled more accurately with prolonged periods of elevated antigenic stimulation or replicative stress, rather than as discrete spikes (as in the case of acute infections). Nuanced stress paradigms could also be introduced to capture the wide range of psychological, physiological, and pathological stressors that individuals experience over a lifetime.

Looking ahead, longitudinal human studies stand out as a critical avenue for gathering the empirical data needed to validate, refine, or revise the theoretical constructs outlined here. By following cohorts over extended periods—spanning decades if possible—investigators could track how the balance of naïve and memory T cells evolves in relation to infection history, stress levels (assessed through questionnaires, biomarkers like cortisol, or social-environmental indices), and emerging clinical signs of autoimmune dysregulation. Such data would enable researchers to calibrate model parameters more precisely, to identify which patterns of infection or stress are most strongly correlated with precipitous declines in naïve cells, and to uncover early immunological indicators of impending autoimmunity.

The potential translational impact of this research lies in formulating interventions that mitigate these detrimental shifts in T cell homeostasis. Strategies might include targeted vaccination programs aimed at reducing the incidence or severity of high-burden infections that disproportionately drive memory inflation. Stress reduction techniques, mental health support, and lifestyle interventions could also help preserve immune regulatory capacity, potentially slowing the skew toward memory-dominant profiles. Moreover, novel therapies to modulate or rejuvenate thymic output—ranging from hormonal treatments to regenerative medicine approaches—might offer ways to sustain a healthier balance in older adults. In the future, therapies designed to remove or reprogram senescent memory T cells could also be explored, potentially reducing the inflammatory burden that fuels tissue damage and autoimmune flares.

Still, the question of causality versus correlation remains a challenging frontier. Although this article and its associated model posit that repeated infections and stress likely accelerate the natural age-driven inversion of T cell profiles, many confounding factors must be disentangled. Genetic predisposition, environmental toxins, nutritional status, the gut microbiome, and coexisting chronic diseases (like diabetes or cardiovascular conditions) all modulate immune function and may act in synergy with or independently of infection and stress burdens. Dissecting these interacting influences will require a systems immunology approach, combining big-data analytics, multi-omics profiling (transcriptomics, proteomics, metabolomics), and carefully designed longitudinal sampling.

Nonetheless, the overall conclusion remains compelling: the shift from a naïve- rich to a memory-rich T cell compartment is a hallmark of aging, and repeated immunological assaults, together with episodes of stress, appear to drive this process more aggressively. Although memory T cells confer protective advantages against previously encountered pathogens, an imbalance wherein naïve cells become scarce can leave the immune system less flexible and more prone to autoreactivity. By recognizing and modeling these dynamics, we open pathways to not only predict the course of immunosenescence but also to intervene in ways that could preserve immune resilience. Such insights have broad implications for public health, geriatric medicine, autoimmune disease management, and future vaccine or therapeutic development.

In conclusion, the data and modeling framework discussed here highlight the delicate equilibrium underlying T cell homeostasis and the ways in which aging, infection history, and stress intersect to potentially reshape immunological risk profiles. Further refinement of the proposed equations, along with the integration of deeper biological detail, promises to shed even more light on the pathways leading to autoimmune susceptibility. As aging populations grow and global infection patterns evolve, understanding how to preserve a healthy, functionally broad T cell repertoire could play a key role in minimizing disease burden and improving quality of life in later years. By drawing upon a mix of computational modeling, clinical observation, and mechanistic immunology, researchers and clinicians alike can work toward a future in which the detrimental effects of immunosenescence are mitigated—leading to healthier aging and reduced prevalence of autoimmune dysfunction.

*The Author claims that there are no conflicts of interest.

## Section 6. Attachment

### Python Code

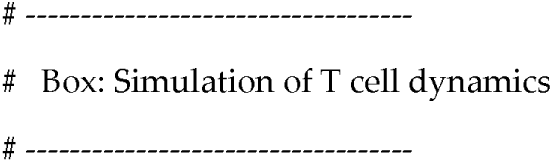
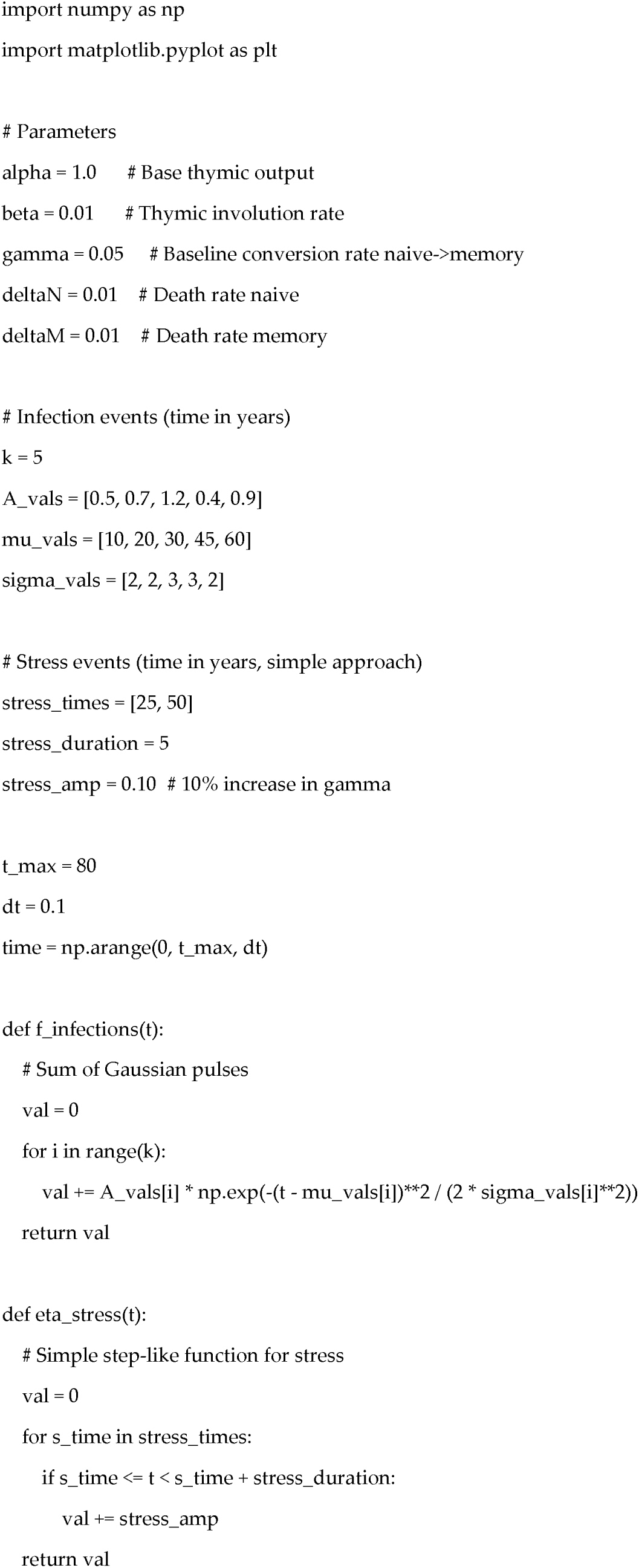
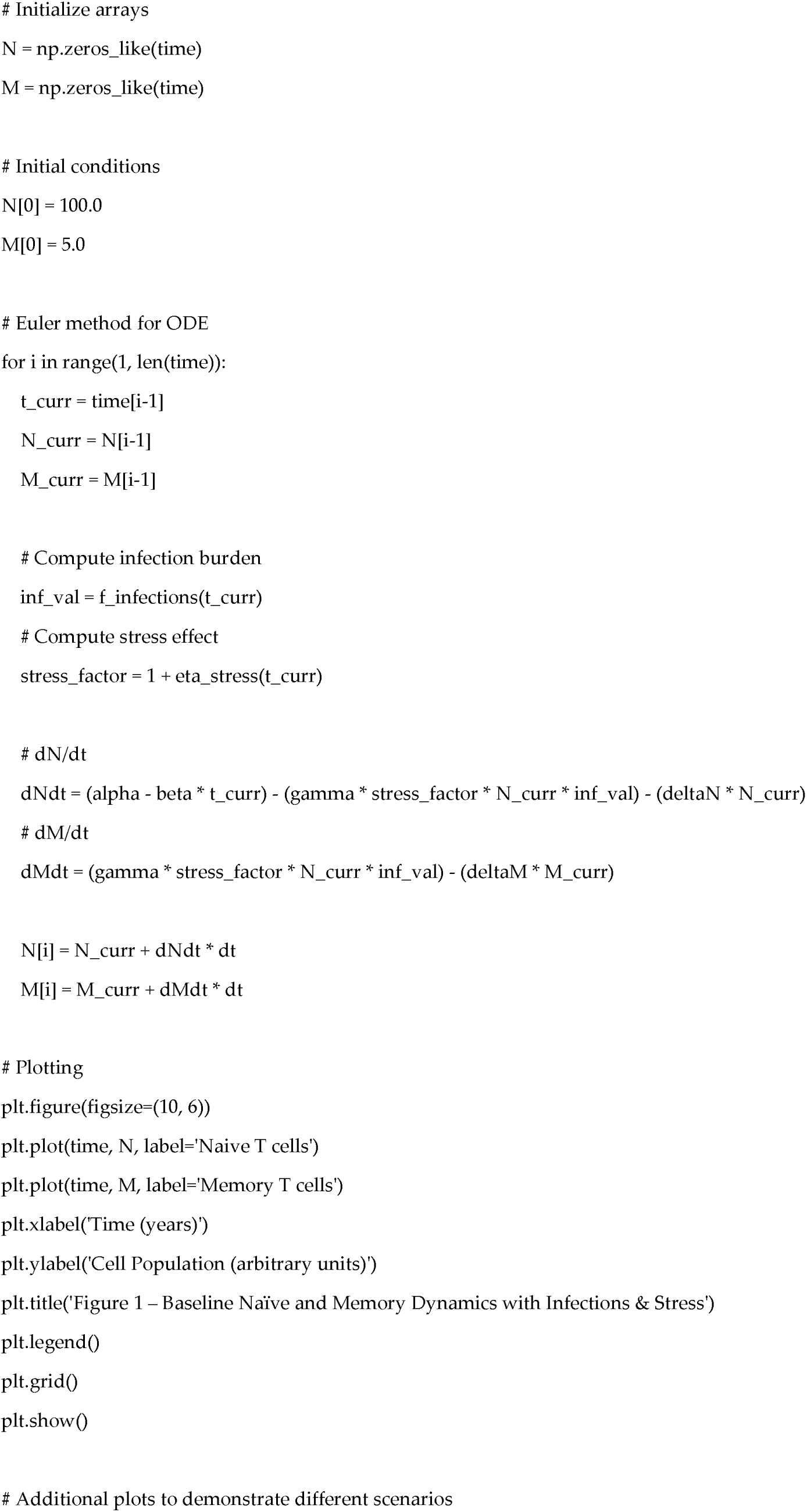
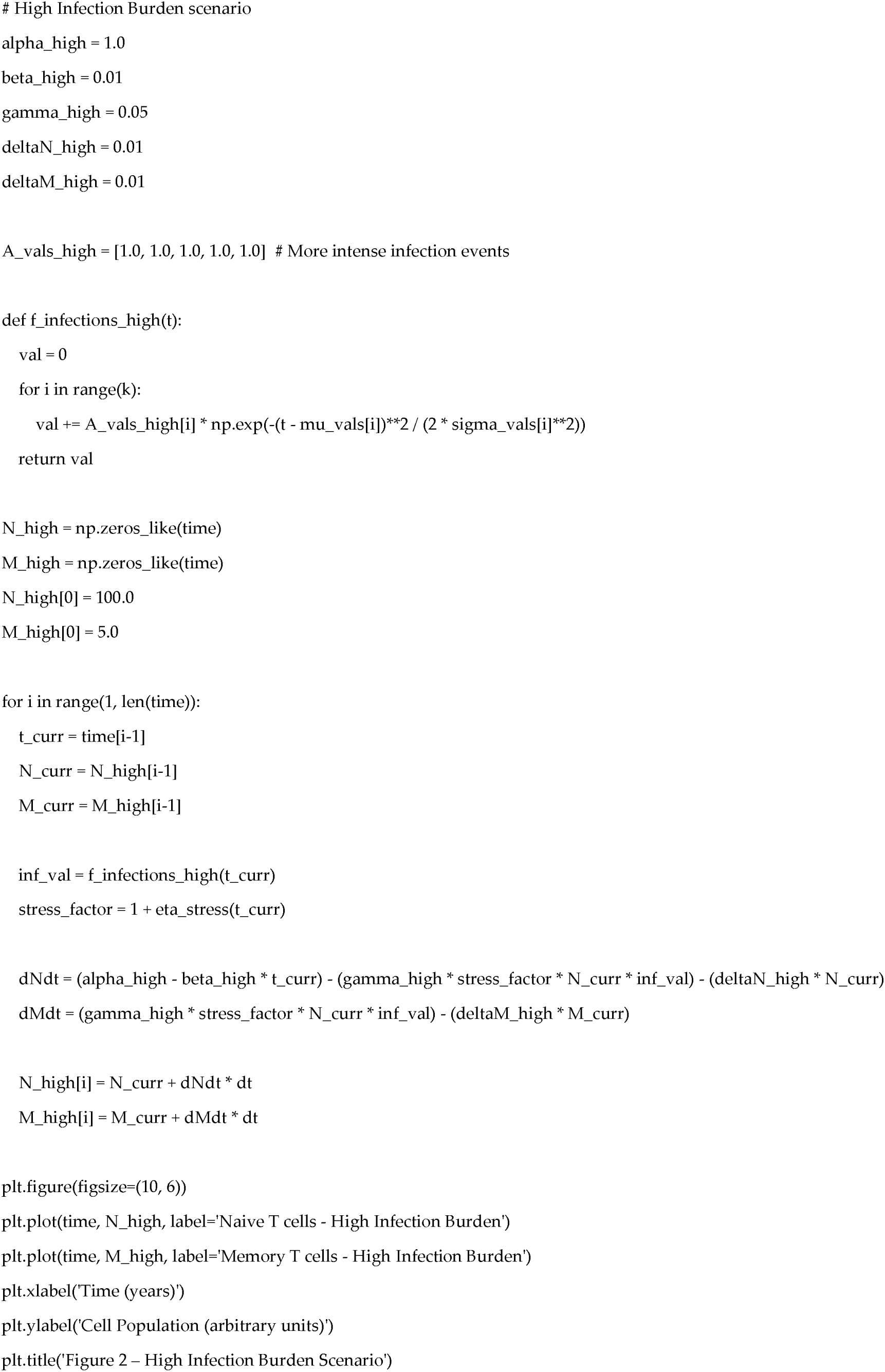
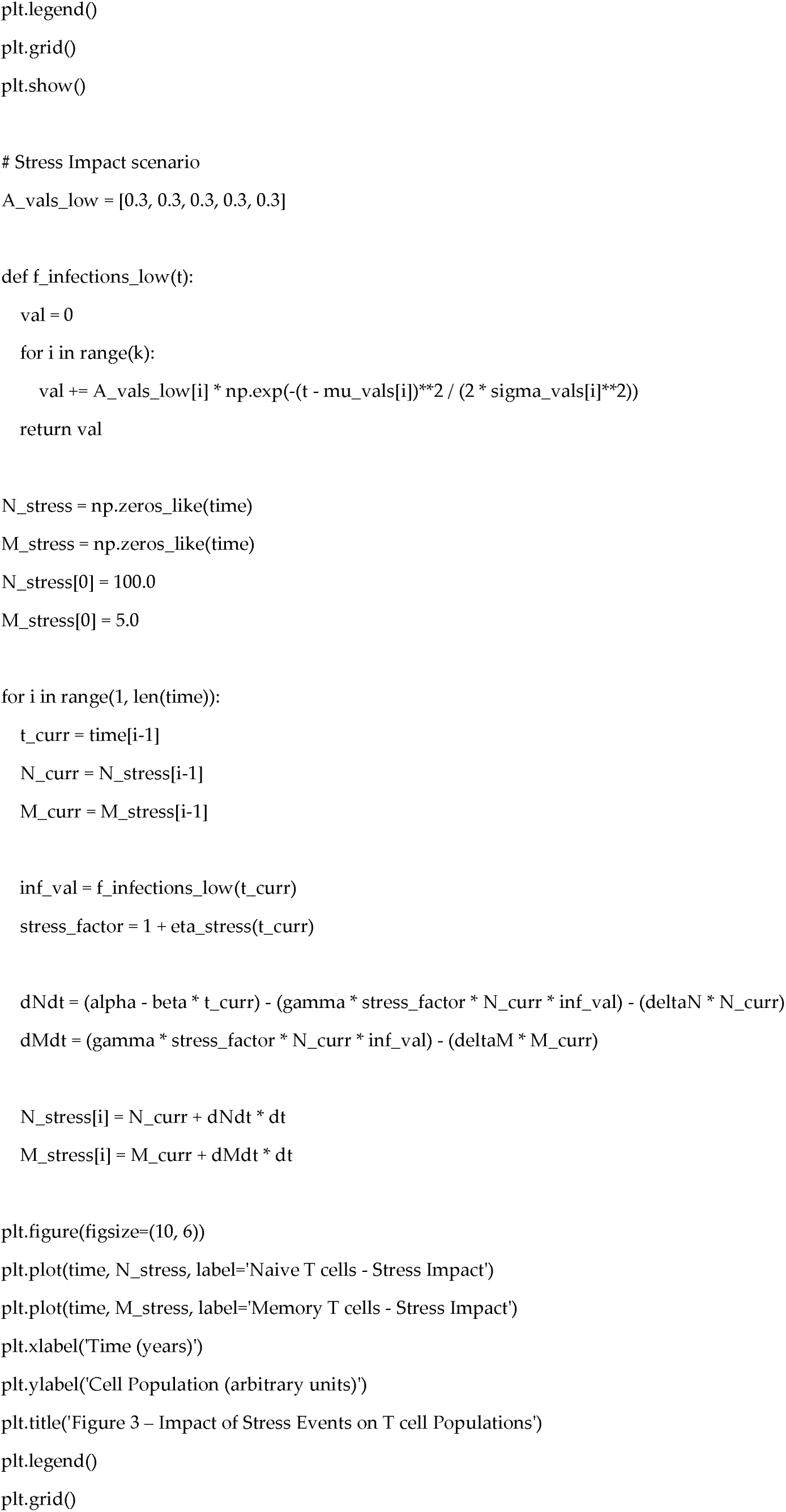
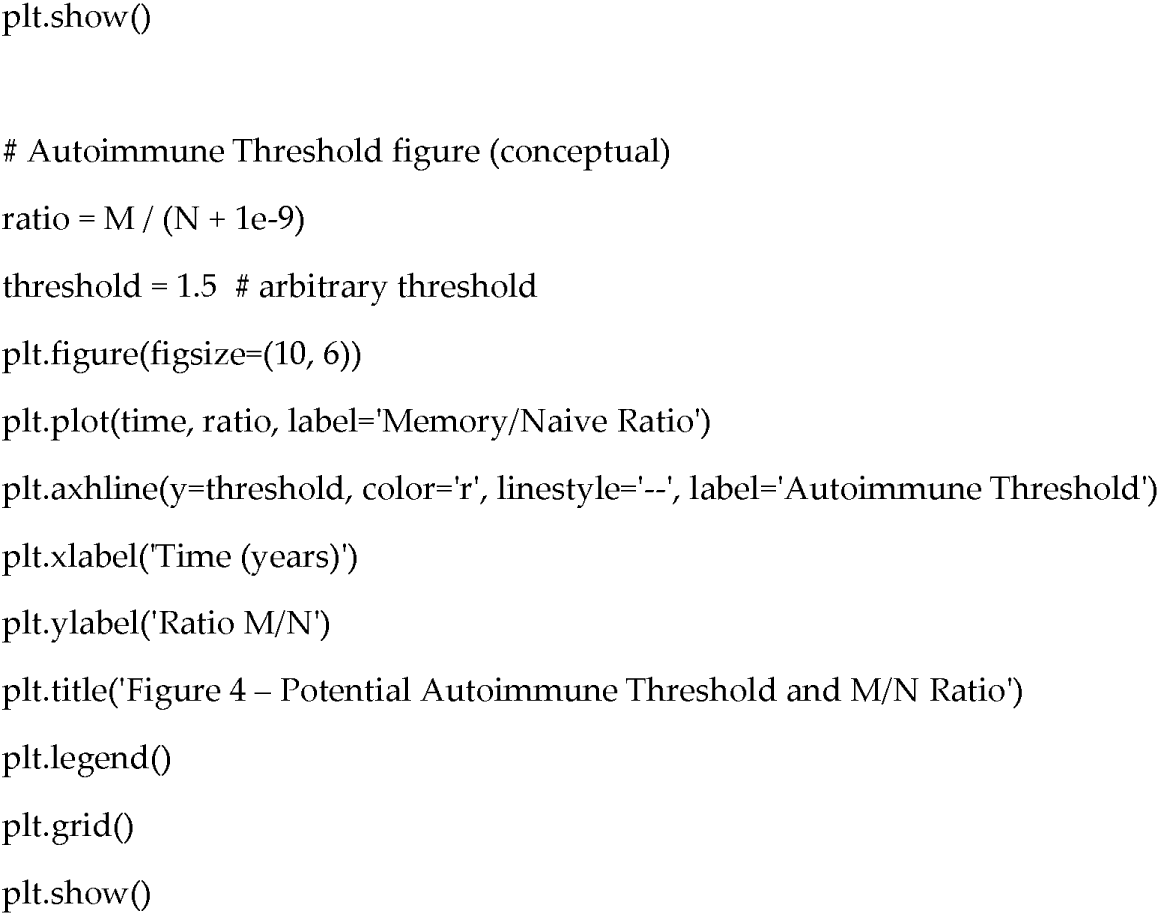

## Notes

### Competing Interest Statement

The authors have declared no competing interest.

## Section 7. References (Alphabetical Order)

1. Abbas A.K., Lichtman A.H., Pillai S. (2018). Cellular and Molecular Immunology. 9th ed. Philadelphia: Elsevier.

2. Amital H., Shoenfeld Y., Amital D., et al. (2008). “Stress and Autoimmunity: New Insights.” Autoimmunity Reviews, 7(3), 214–217.

3. Anderson R.M., May R.M. (1991). Infectious Diseases of Humans: Dynamics and Control. Oxford: Oxford University Press.

4. Britanova O.V., Putintseva E.V., Shugay M., et al. (2016). “Dynamics of Individual T Cell Repertoires: From Cord Blood to Centenarians.” Journal of Immunology, 196(12), 5005–5013.

5. Carr E.J., Dooley J., Garcia-Perez J.E., et al. (2022). “The Impact of Age and Genetic Variation on T Cell Function.” Nature Immunology, 23, 584–595.

6. Dawes R., Petrovas C., O’Connor M., et al. (2019). “T Cell Receptor Repertoire Divergence in Autoimmunity and Chronic Infections.” Frontiers in Immunology, 10, 237.

7. Franceschi C., Garagnani P., Parini P., et al. (2018). “Inflammaging: A New Immune–Metabolic Viewpoint for Age-Related Diseases.” Nature Reviews Endocrinology, 14(10), 576–590.

8. Fulop T., Larbi A., Khalil A., et al. (2018). “Are We Ill Because We Age? Frontiers in Immunosenescence.” Biogerontology, 19(6), 497–510.

9. Glaser R., Kiecolt-Glaser J.K. (2005). “Stress-Induced Immune Dysfunction: Implications for Health.” Nature Reviews Immunology, 5(3), 243–251.

10. Karrer U., Sierro S., Wagner M., et al. (2003). “Memory Inflation: Continuous Accumulation of Antiviral CD8+^++ T Cells Over Time.” Journal of Immunology, 170(4), 2022–2029.

11. Montgomery. R. M. (2024). “Molecular Mechanisms of Chronic Stress in Immune Dysregulation: From Cytokine Networks to Clinical Manifestations”. DOI: 10.62162/WNSC10609.2.

12. Montgomery R. M. (2024)a. *“*Frailty Syndrome A Global Perspective on Prevalence, Challenges, and Strategies for Prevention and Management*”.* DOI: 10.20944/preprints202409.1249.v1.

13. Murphy K., Weaver C. (2017). Janeway’s Immunobiology. 9th ed. New York: Garland Science.

14. Nikolich-Žugich J. (2018). “The Twilight of Immunity: Emerging Concepts in Immunosenescence.” Nature Immunology, 19(1), 10–19.

15. Zhang H., Weyand C.M., Goronzy J.J. (2019). “T Cell Ageing and the Links to Chronic Inflammation.” Nature Reviews Rheumatology, 16(2), 87–101.

